# Muscle activation patterns are more constrained and regular in treadmill than in overground human locomotion

**DOI:** 10.1101/2020.07.07.191080

**Authors:** Ilaria Mileti, Aurora Serra, Nerses Wolf, Victor Munoz-Martel, Antonis Ekizos, Eduardo Palermo, Adamantios Arampatzis, Alessandro Santuz

**Affiliations:** Department of Mechanical and Aerospace Engineering, Sapienza University of Rome, 00185 Rome, Italy; Department of Electrical Engineering and Informatics, Technische Universität Berlin, 10587 Berlin, Germany; Department of Training and Movement Sciences, Humboldt-Universität zu Berlin, 10115 Berlin, Germany; Berlin School of Movement Science, Humboldt-Universität zu Berlin, 10115 Berlin, Germany; Atlantic Mobility Action Project, Brain Repair Centre, Department of Medical Neuroscience, Dalhousie University, Halifax, Nova Scotia B3H 4R2, Canada

**Keywords:** Locomotion, muscle synergies, motor control, treadmill locomotion, overground locomotion

## Abstract

The use of motorized treadmills as convenient tools for the study of locomotion has been in vogue for many decades. However, despite the widespread presence of these devices in many scientific and clinical environments, a full consensus on their validity to faithfully substitute free overground locomotion is still missing. Specifically, little information is available on whether and how the neural control of movement is affected when humans walk and run on a treadmill as compared to overground. Here, we made use of linear and nonlinear analysis tools to extract information from electromyographic recordings during walking and running overground, and on an instrumented treadmill. We extracted synergistic activation patterns from the muscles of the lower limb via non-negative matrix factorization. We then investigated how the motor modules (or time-invariant muscle weightings) were used in the two locomotion environments. Subsequently, we examined the timing of motor primitives (or time-dependent coefficients of muscle synergies) by calculating their duration, the time of main activation, and their Hurst exponent, a nonlinear metric derived from fractal analysis. We found that motor modules were not influenced by the locomotion environment, while motor primitives resulted overall more regular in treadmill than in overground locomotion, with the main activity of the primitive for propulsion shifted earlier in time. Our results suggest that the spatial and sensory constraints imposed by the treadmill environment forced the central nervous system to adopt a different neural control strategy than that used for free overground locomotion. A data-driven indication that treadmills induce perturbations to the neural control of locomotion.

## Introduction

Amongst the various behaviors that can be used to investigate the neural control of movement, locomotion is an ideal choice: automatized, synergistic, general, cyclic, and phylogenetically old, it embodies many scientifically convenient characteristics (Bernstein, 1967). However, the study of overground locomotion in free, open spaces is often unfeasible due to logistical, technological, and other limitations. Motorized treadmills are an intuitive solution to simplify the analysis of locomotion and are nowadays of widespread use in research, clinical practice and sports-related training (Miller et al., 2019; Van Hooren et al., 2019). Nevertheless, despite their broad use, a full consensus as to whether treadmills are a valid means to generalize findings on the neural control of free locomotion is yet to be found (Rozumalski et al., 2015; Oliveira et al., 2016; Miller et al., 2019; Van Hooren et al., 2019).

Treadmill locomotion is often considered an invalid alternative to overground locomotion due to the mechanical advantage introduced by the moving belt. However, already in 1980, the Dutch biomechanist van Ingen Schenau showed that there are no mechanical differences between treadmill and overground locomotion as long as the belt’s speed remains constant (van Ingen Schenau, 1980). Yet, other factors might affect the physiological determinants of treadmill walking and running: the compliance of the surface, the lack of air resistance, the fixed rather than moving visual feedback, the degree of habituation, among others. (Jones and Doust, 1996; Parvataneni et al., 2009; Mooses et al., 2014; Miller et al., 2019; Van Hooren et al., 2019). For instance, when comparing the energetics and performance outcomes of treadmill and overground running in humans, a great variability across studies arises, some of which is related to the different speeds used for the investigation (Miller et al., 2019). The kinematics and kinetics of running seem to be largely independent on the chosen locomotion environment (Van Hooren et al., 2019). As for walking, kinematics and kinetics can vary between treadmill and overground (Hollman et al., 2016; Yang and King, 2016), but studies on the behavior of the triceps surae muscle fascicles and electromyographic (EMG) activity of the lower limbs did not find significant differences (Cronin and Finni, 2013; Ibala et al., 2019). Generally speaking, there is widespread scientific proof that the kinematics, kinetics, and EMG activity recorded during treadmill and overground locomotion are similar enough to allow the use of treadmill for scientific purposes (Lee and Hidler, 2008; Riley et al., 2008; Parvataneni et al., 2009; Chia et al., 2014).

In this study, we set out to investigate the modular organization of muscle activity during overground and treadmill walking and running. We started from the general hypothesis that previously found similarities in kinematics, kinetics and EMG activity do not necessarily imply that locomotion in different environments is controlled with similar neuromotor strategies. Since it is known that when movement is constrained by internal or external factors the neuromotor control is affected (Martino et al., 2015; Santuz et al., 2019, 2020a), we sought to put together a new set of analysis tools designed to be more sensitive to such variations. Thus, to better understand the neural control processes underlying locomotion in different environments, we adopted a novel framework based on both linear and nonlinear approaches for extracting information from EMG data. First, we used non-negative matrix factorization (NMF) as a linear decomposition tool to extract muscle synergies from the EMG activity recorded from the lower limb during walking and running (Bernstein, 1967; Bizzi et al., 1991, 2008; Lee and Seung, 1999; Santuz et al., 2017a). Then, we analyzed the motor modules, or the weighted contributions of each muscle activity, and the timing characteristics of motor primitives, which are the time-dependent components of muscle synergies (Santuz et al., 2018a). Lastly, we used fractal analysis to compute the Hurst exponent of motor primitives, in order to gain deeper insight into their temporal structure (Santuz and Akay, 2020). By using these tools, we recently found that both internal and external perturbations applied to human and murine locomotion affect the timing of motor primitives, despite minor changes in the number and composition of motor modules (Santuz et al., 2018b, 2019, 2020a, 2020b; Santuz and Akay, 2020). Specifically, we could systematically associate a relatively longer duration of motor primitives (i.e. increased width of the signal) in genetically modified mice lacking proprioceptive feedback from muscle spindles (Santuz et al., 2019), in aging humans as compared to young (Santuz et al., 2020a), and in young adults walking and running on uneven terrain (Santuz et al., 2018b), on unstable ground (Santuz et al., 2020a) or running at extremely high speeds (Santuz et al., 2020b).

Here, we aimed at uncovering some neuromotor control features of overground and treadmill locomotion using a novel combination of machine learning and fractal analysis. Based on our previous findings on perturbed and unperturbed locomotion (Santuz et al., 2018b, 2019, 2020a; Santuz and Akay, 2020), we hypothesized that: a) treadmill, as compared to overground, would perturb more the locomotor system due to the increased sensory and spatial constraints; and b) the neural control of treadmill locomotion would be forced to be more regular by the aforementioned constraints, thus resulting in motor primitives having a Hurst exponent lower in treadmill than in overground.

## Results

### Gait parameters

The stance and swing times and the cadence are reported in Figure 1. The coefficient of variation of stance, swing and cadence, together with the strike index are reported in Table 1. A main effect of locomotion type (walking compared to running) was found for stance, swing and cadence (p < 0.001). Stance and swing phase duration were significantly lower and cadence higher in running (p < 0.001). Treadmill, compared to overground locomotion, made swing times decrease (p = 0.032). No environment by type interaction (p > 0.05) was observed for any of the gait parameters. All the other comparisons were statistically (p > 0.05) not significant.

**Table 1.** The step-to-step percent coefficient of variation (CV) of stance, swing and cadence is reported as the ratio between mean and standard deviation. The strike index is the distance of the center of pressure at touchdown from the most posterior part of the heel, normalized to foot length.

| Locomotion type | Environment | Stance CV | Swing CV | Cadence CV | Strike index |
| --- | --- | --- | --- | --- | --- |
| Walking | Overground | $3.8 \pm 3.5\%$ | $4.4 \pm 4.2\%$ | $2.1 \pm 2.1\%$ | $0.063 \pm 0.030$ |
| | Treadmill | $2.5 \pm 1.8\%$ | $4.0 \pm 3.0\%$ | $1.2 \pm 0.4\%$ | $0.047 \pm 0.018$ |
| Running | Overground | $7.4 \pm 3.4\%$ | $4.3 \pm 1.7\%$ | $2.6 \pm 1.4\%$ | $0.066 \pm 0.029$ |
| | Treadmill | $7.1 \pm 15.9\%$ | $4.5 \pm 8.6\%$ | $1.9 \pm 2.3\%$ | $0.073 \pm 0.051$ |

**Figure 1.**
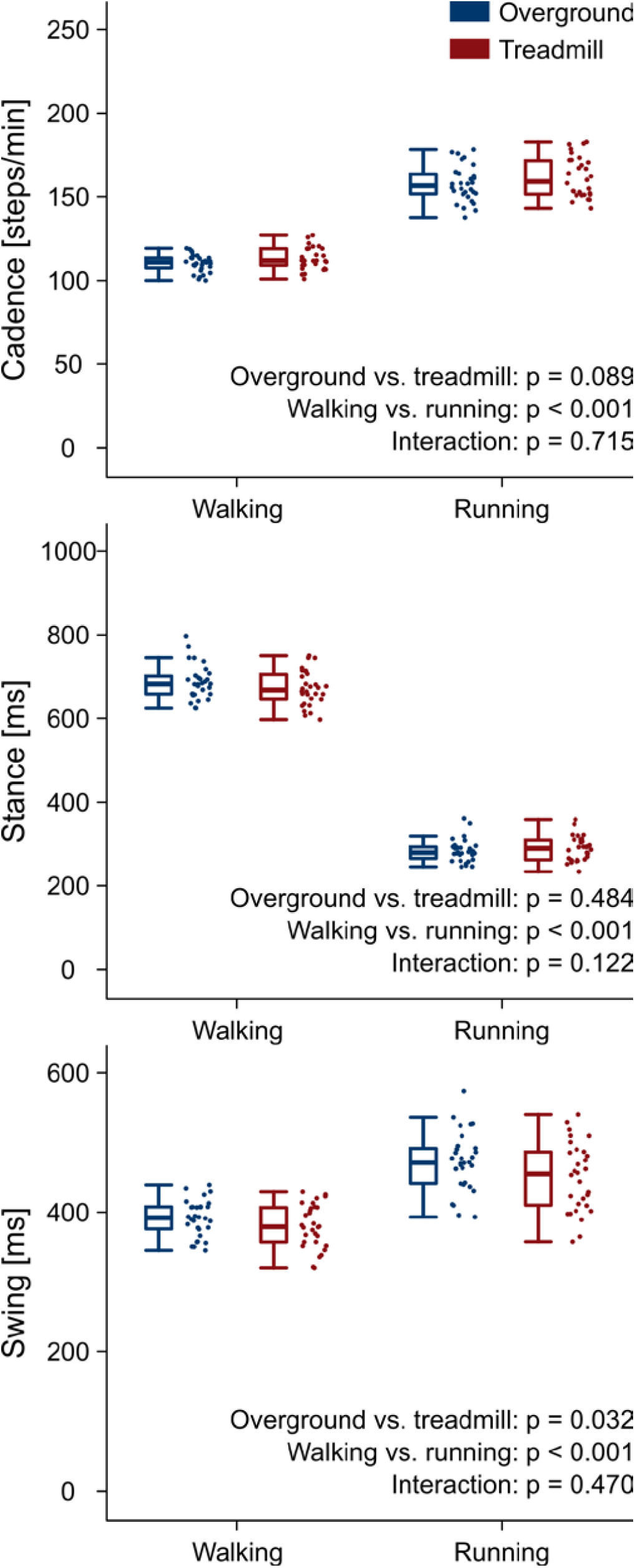
Gait parameters. Boxplots depicting stance, swing, and cadence for the four locomotion conditions.

### Muscle synergies

The minimum number of synergies necessary to reconstruct the EMG data (i.e. the NMF factorization rank) was 4.6 ± 0.5 for overground walking, 4.5 ± 0.6 for treadmill walking, 4.2 ± 0.6 for overground running and 4.6 ± 0.7 for treadmill running, with no significant differences (overground vs. treadmill p = 0.077; walking vs. running p = 0.239). The average reconstruction quality (i.e. the R^2^ or the EMG variability accounted for by the factorization) was 0.829 ± 0.028 for overground walking, 0.843 ± 0.027 for treadmill walking, 0.850 ± 0.025 for overground running and 0.869 ± 0.026 for treadmill running. An effect of locomotion environment (overground vs. treadmill, p = 0.001) and type (walking vs. running, p < 0.001) was found for the reconstruction quality, but no environment by type interactions (p = 0.567). The percentage of combined synergies was 16.1% for overground walking, 19.1% for treadmill walking, 19.0% for overground running and 23.0% for treadmill running.

Four fundamental synergies were clustered in all gait conditions. In both walking (Figure 2) and running (Figure 3), the first synergy functionally referred to the body weight acceptance, with a major involvement of knee extensors and hip extensors and abductors. The second synergy described the propulsion phase, to which the plantarflexors mainly contributed. The third synergy identified the early swing, showing the involvement of foot dorsiflexors. The fourth and last synergy reflected the late swing and the landing preparation, highlighting the relevant influence of knee flexors (in both walking and running), and foot dorsiflexors (mostly in running). No main effect of the locomotion environment was found for any of the motor modules in walking or running. In walking, the SPM analysis detected significant differences in the descending part of the late swing primitive (between the 179^th^ and 185^th^ normalized time points, p = 0.001), as shown in Figure 2. In running (Figure 3), the SPM highlighted differences in both the ascending (points 34 to 44, p < 0.001) and descending (points 63 to 82, p < 0.001) portion of the propulsion primitive.

**Figure 2.**
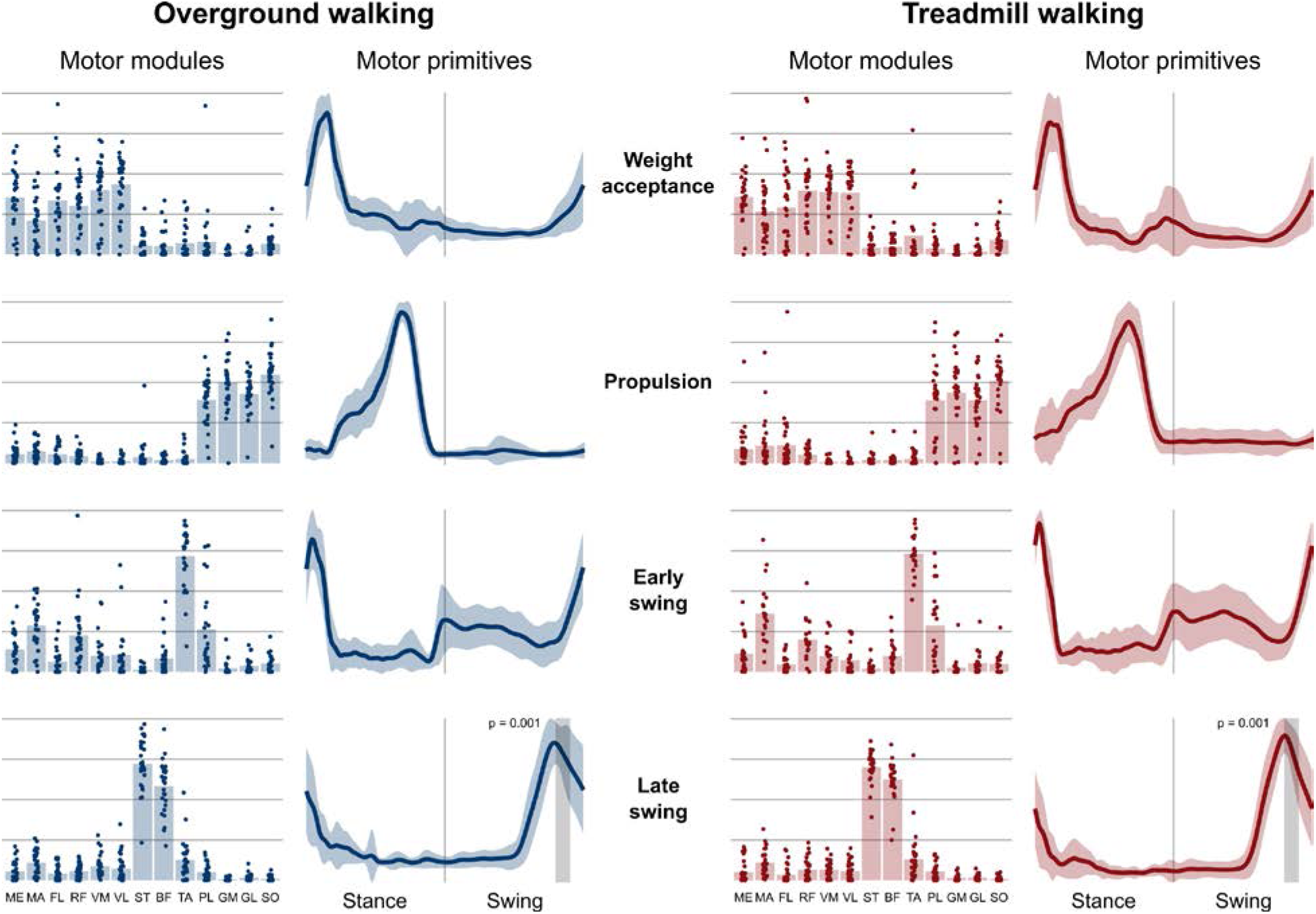
Motor modules and motor primitives of the four fundamental synergies for overground and treamill walking. The motor modules are presented on a normalized y-axis base: each muscle contribution within one synergy can range from 0 to 1 and each point represents individual trials. For the mean motor primitives (shaded standard deviation), the x-axis full scale represents the averaged gait cycle (with stance and swing normalized to the same amount of points and divided by a vertical line) and the y-axis the normalized amplitude. Differences in motor primitives between overground and treadmill found by statistical parametric mapping are shown as vertical shaded areas with relevant p-value. Muscle abbreviations: ME=gluteus medius, MA=gluteus maximus, FL=tensor fasciæ latæ, RF=rectus femoris, VM=vastus medialis, VL=vastus lateralis, ST=semitendinosus, BF=biceps femoris, TA=tibialis anterior, PL=peroneus longus, GM=gastrocnemius medialis, GL=gastrocnemius lateralis, SO=soleus.

**Figure 3.**
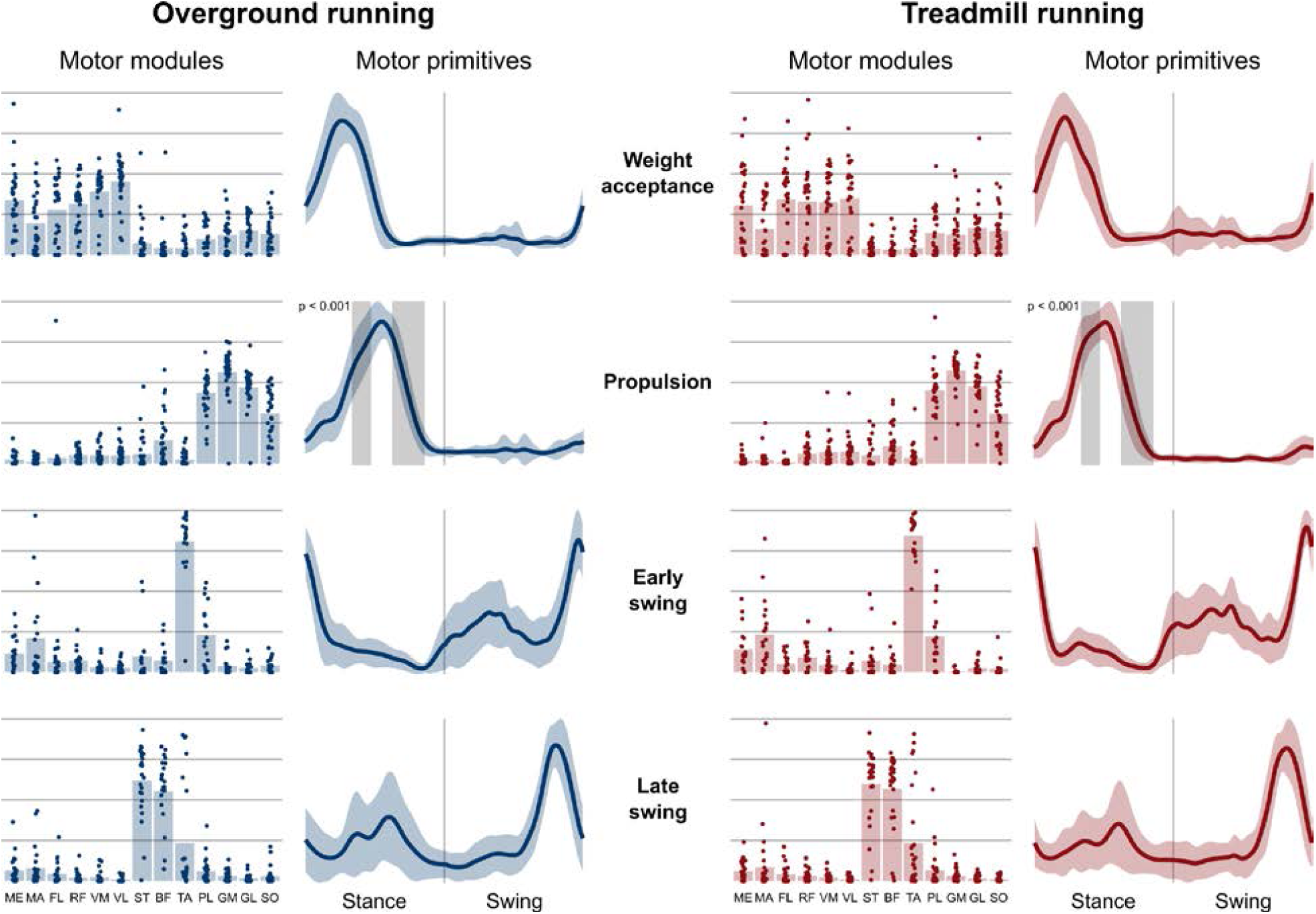
Motor modules and motor primitives of the four fundamental synergies for overground and treamill running. The motor modules are presented on a normalized y-axis base: each muscle contribution within one synergy can range from 0 to 1 and each point represents individual trials. For the mean motor primitives (shaded standard deviation), the x-axis full scale represents the averaged gait cycle (with stance and swing normalized to the same amount of points and divided by a vertical line) and the y-axis the normalized amplitude. Differences in motor primitives between overground and treadmill found by statistical parametric mapping are shown as vertical shaded areas with relevant p-value. Muscle abbreviations: ME=gluteus medius, MA=gluteus maximus, FL=tensor fasciæ latæ, RF=rectus femoris, VM=vastus medialis, VL=vastus lateralis, ST=semitendinosus, BF=biceps femoris, TA=tibialis anterior, PL=peroneus longus, GM=gastrocnemius medialis, GL=gastrocnemius lateralis, SO=soleus.

### Motor primitive geometrics

The CoA of the propulsion primitive shifted earlier in time when switching from overground to treadmill locomotion in both walking and running (Figure 4). Moreover, the CoA of motor primitives was different between walking and running in all synergies: higher in weight acceptance and early swing; lower in propulsion and late swing (Figure 4). The weight acceptance and propulsion primitives were wider (i.e. higher FWHM) relative to the stance phase in running than in walking, but the locomotion environment did not show an effect on FWHM. The widening is visible in both the box plots of Figure 5 and the heat maps of Figure 6.

**Figure 4.**
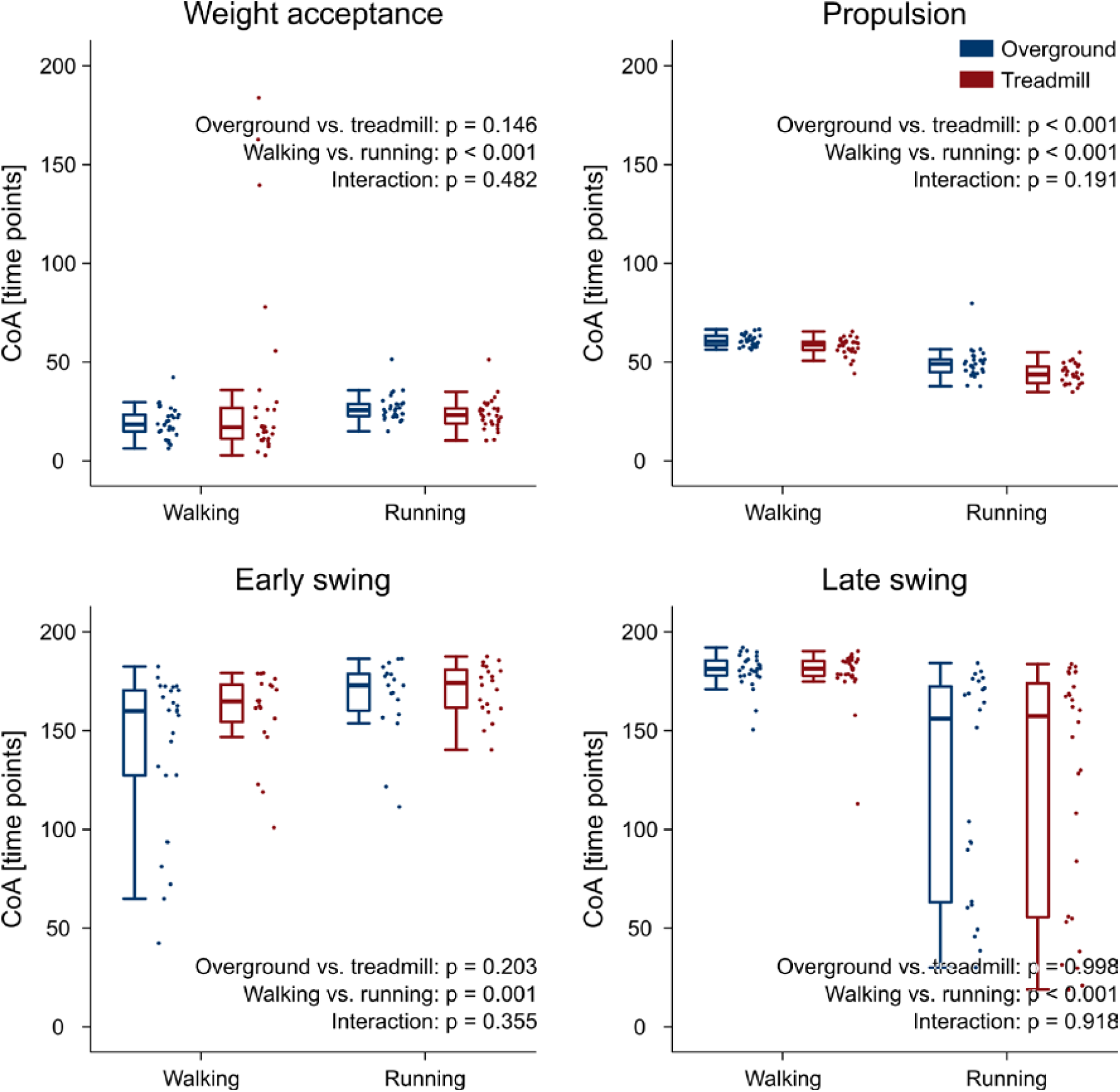
Box plots representing the center of activity (CoA) values for the motor primitives of the four fundamental muscle synergies. Individual trial values are presented as points.

**Figure 5.**
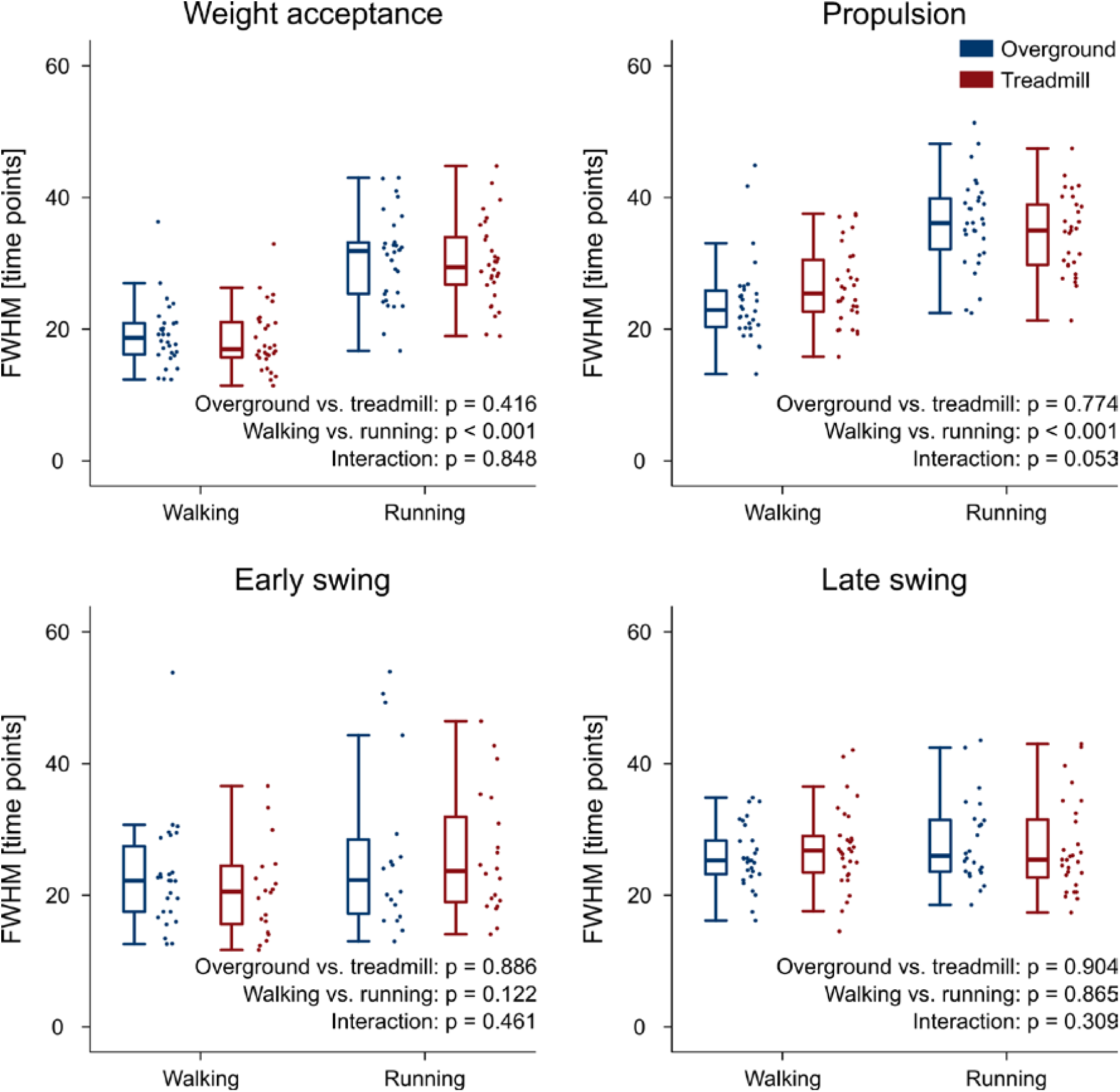
Box plots representing the full width at half maximum (FWHM) values for the motor primitives of the four fundamental muscle synergies. Individual trial values are presented as points.

**Figure 6.**
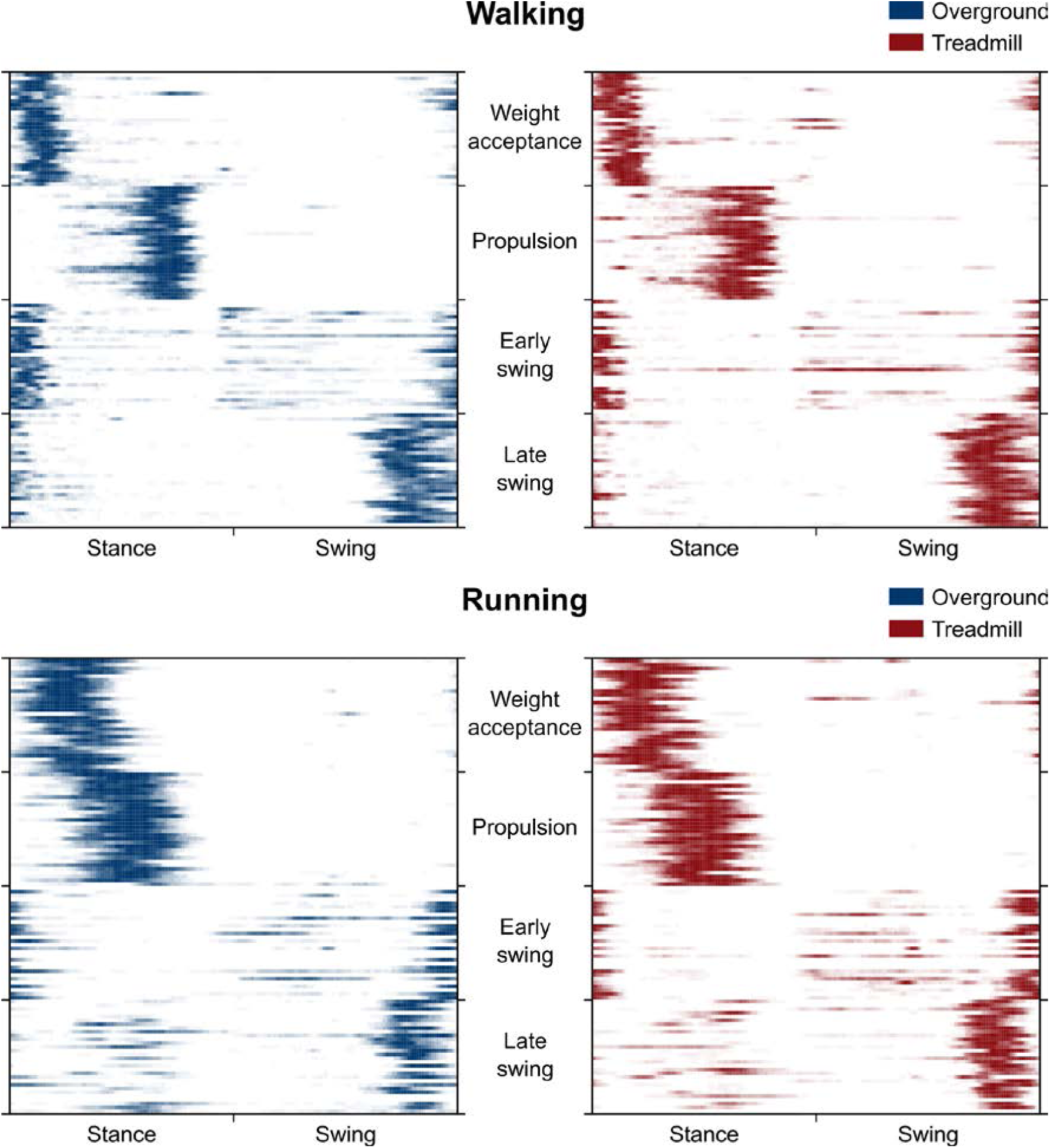
Heat maps representing the average occurrence of values bigger than half maximum for each trial (rows of the maps). We calculated trial-by-trial, for each of the 200 time points (columns of the maps) and gait cycle, the number of times a motor primitive was exceeding half maximum and reported the mean results in a color-coded fashion: from white (the primitive never exceeded half maximum) to blue or red (the primitive exceeded half maximum in all the 30 gait cycles of that trial). Missing primitives are reported as fully white rows.

### Fractal analysis of motor primitives

The H values and the rescaled range versus window length log-log plots are shown in Figure 7. H values of motor primitives were lower in treadmill compared to overground in both walking and running. Moreover, the mean H values were lower than 0.5 in all four conditions, indicating anti-persistent behavior of motor primitives (Mandelbrot, 1983; Gneiting and Schlather, 2004). Anti-persistence means that, in the motor primitives of treadmill locomotion, short-term oscillations between high and low values were less random than in overground. In other words, the power-like decay of motor primitive’s autocorrelation was faster in treadmill than in overground locomotion (Tarnopolski, 2016).

**Figure 7.**
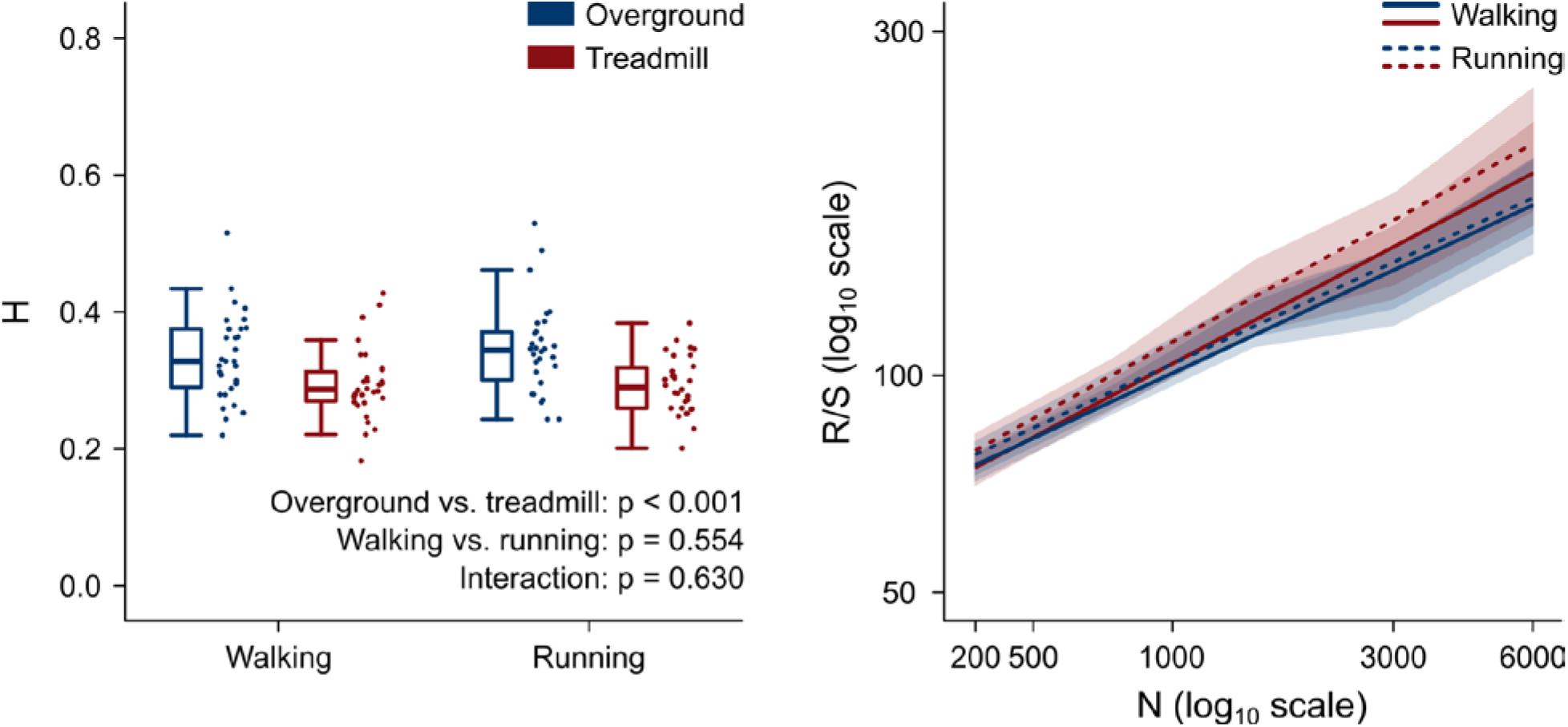
Left: box plots representing the Hurst exponent (H) values, calculated as the average exponent of all primitives per trial. Individual trial values are presented as points. Right: log-log plot of the rescaled range (R/S) versus window size (n, in number of normalized time points) for the four locomotion conditions. The slope of each regression line is H. Standard deviations are presented as shaded areas around each relevant regression line.

## Discussion

Fractal analysis revealed that motor primitives were more regular in treadmill than in overground locomotion, as hypothesized. This novel finding suggests that the spatial and sensory constraints imposed by the treadmill environment forced the CNS to adopt a different neural control strategy. While no difference was found in the FWHM of motor primitives for overground and treadmill locomotion, we could show that the CoA of the propulsion primitive was shifted earlier in time when our participants walked and ran on a treadmill. This partially confirmed our hypothesis that treadmills induce perturbations to locomotion.

Gait spatiotemporal parameters were in general scarcely affected by the locomotion environment. We found only a decreased swing duration in treadmill compared to overground running. This is in agreement with most of the relevant literature reports (Lee and Hidler, 2008; Parvataneni et al., 2009; Oliveira et al., 2016; Van Hooren et al., 2019; Santuz et al., 2020a). Moreover, the coefficient of variability of the cadence, measured in steps per minute, was significantly lower in treadmill for both walking and running, suggesting a higher degree of regularity of treadmill locomotion imposed by the less variable speed (Dingwell and Cusumano, 2000).

We and others showed in previous studies that both the number and function of muscle synergies are largely shared across locomotion types and settings. For instance, in mice the number of synergies for walking and swimming is identical (Santuz et al., 2019) as it is in humans for walking and running (Cappellini et al., 2006; Lacquaniti et al., 2012; Santuz et al., 2017a) or in locomotion at different speeds (Ivanenko et al., 2004; Santuz et al., 2020b). When adding external (e.g. mechanical) or internal (e.g. aging or pathology) perturbations to locomotion, the number of synergies is not affected in both mice (Santuz et al., 2019) and humans (Maclellan et al., 2014; Santuz et al., 2018b, 2020a; Holubarsch et al., 2019; Janshen et al., 2020). Several studies have attempted in the past to highlight potential discrepancies between overground and treadmill locomotion from many perspectives concluding that spatiotemporal, kinematic, kinetic, and muscle-tendon interaction measures are scarcely influenced by the locomotion environment (Van Hooren et al., 2019). One study also examined muscle synergies, finding that motor primitives underwent “minimal temporal adjustments” (Oliveira et al., 2016). Here, we extended that investigation by adding walking to the analysis, unmistakably confirming that the number of muscle synergies was conserved across locomotion types (i.e. walking and running) and environments (i.e. overground and treadmill). The four extracted synergies described the two macro phases of the gait cycle: the stance (weight acceptance and propulsion synergies) and the swing (early and late swing synergies), similar to what was found in other studies (Santuz et al., 2018b, 2018c, 2020a, 2020b). These observations indicate that overground and treadmill locomotion share largely similar modular organization of muscle activations, despite small temporal adjustments of motor primitives. A fact, however, that does not exclude further alterations of the neuromotor control that are invisible to the naked eye.

Fractal analysis can expose local or global properties of a time series that would be otherwise hardly visible to the naked eye and/or simply too difficult to quantify (Santuz and Akay, 2020). In a recent study where we used the Higuchi’s fractal dimension, we found similar local complexity of motor primitives for overground and treadmill locomotion (Gneiting and Schlather, 2004; Santuz et al., 2020a). Here however, we set out to analyze the global, rather than local, fractal properties of motor primitives by estimating the parameter H. There is no specific advantage in analyzing the local or global fractal properties of motor primitives: both characteristics are informative, even though on a different level. While the Higuchi’s fractal dimension tells us what happens in the short term by “zooming in” on the signal, H helps us depicting a general picture of what happens in the long term (Gneiting and Schlather, 2004; Santuz and Akay, 2020). H can vary between 0 and 1, with H = 0.5 indicating a random series (Mandelbrot, 1983; Qian and Rasheed, 2004). For 0.5 < H < 1, in the long term a positive or negative trend is visible, making the time series persistent or with “long memory” (Mandelbrot, 1983; Gneiting and Schlather, 2004). For 0 < H < 0.5, the series is anti-persistent: in the long term high values in the series will be probably followed by low values, with a frequent switch between high and low values as in motor primitives extracted from locomotion (Mandelbrot, 1983; Gneiting and Schlather, 2004). We found that H values were: i) lower than 0.5 in all the analyzed conditions, ii) independent on the locomotion type, and iii) lower in treadmill than in overground locomotion.

First, H < 0.5 is an indication of anti-persistence, meaning that our motor primitives did not show a trend. To make an example of a persistent (i.e. with trend) time series, one can think at the space vs. time graph of a person walking overground at self-paced comfortable speed. Such a curve would be close to a line with slope equal to the speed of the person. As the person walks with almost constant speed, the distance traveled increases as well, showing a positive trend: the distance from the starting point is more likely to increase as time passes rather than to oscillate around a certain value. This example is intuitively dissimilar from the behavior of motor primitives, which are time series that oscillate around a mean value (i.e. they are anti-persistent), due to the fact that locomotion is quasi-periodic (Santuz and Akay, 2020). Thus, it is possible to explain from a physiological perspective why we obtained H < 0.5 for all the analyzed locomotion conditions.

Second, the fact that H is independent on the locomotion type (i.e. walking or running) suggests that speed does not have an influence on the *global* fractal properties of motor primitives. While at increasing locomotion speeds the *local* complexity of motor primitives decreases (Santuz et al., 2020b), the *global* regularity (as measured by H) is not affected. From a neurophysiological point of view, this behavior has no easy explanation. Neural circuits for the control of locomotor type and speed have been found in several regions of the vertebrate CNS: from the diversified populations of inhibitory V1 and excitatory V2a spinal interneurons in the zebrafish spinal cord (Ampatzis et al., 2014; Kimura and Higashijima, 2019), to the human prefrontal cortex (Suzuki et al., 2004; Bulea et al., 2015) and the murine and human brainstem (Al-Yahya et al., 2011; Capelli et al., 2017), passing through the feline cerebellum (Armstrong, 1988) and the V0 and V3 commissural interneurons for left-right alternation and synchronization in the mouse system (Danner et al., 2016). All these circuits have one important thing in common: they implement a flexible modular organization of neuronal excitation and inhibition for smoothly controlling the type and speed of locomotion. Our results suggest that the modular activation of muscles, the final effectors for motion creation and control, is constantly tuned to maintain similar patterns, despite the profound changes happening in the underlying neural circuits.

Third, the found lower H values in treadmill compared to overground suggest that motor primitives for treadmill locomotion are more regular than those for overground walking or running. If primitives were all perfect sinusoidal time series with period equal to the gait cycle, H would be zero. Conversely, if primitives were oscillating around their mean value in a random way, H would be 0.5. It follows that if H decreases from 0.5 to 0, the level of randomness in the time series decreases as well, while regularity increases. A reason for the increased regularity of motor primitives might be the intrinsic regularity of the treadmill belt’s speed (Dingwell and Cusumano, 2000; Riley et al., 2008). Even though it has been shown that treadmill belts slightly decelerate at touchdown only to recover the set speed later in the stance phase and accelerate at lift-off (Van Hooren et al., 2019), the oscillations are likely small and, most importantly, systematic as shown by the lower CV of cadence in both walking and running. The enforced average speed (Dingwell and Cusumano, 2000) and other parameters such as the limited belt dimensions (Van Hooren et al., 2019), could have contributed to make treadmill a more restricted locomotion environment than overground. Physiologically speaking, this suggests that a more regular neural control strategy was needed to overcome the sensory constraints imposed by the treadmill environment, showing that treadmills might be influencing motor coordination more than previously thought. Recently, we used the FWHM of motor primitives as a measure of robustness, concluding that wider (i.e. active for a longer time) primitives indicate more robust motor control in perturbed locomotion settings (Santuz et al., 2018b, 2020a; Janshen et al., 2020). Our idea of robust control is based on the optimal feedback control theory, which postulates that motor systems selectively combine sensory feedback and motor commands to optimize performance (Todorov and Jordan, 2002; Scott, 2004; Tuthill and Azim, 2018). It is known that the treadmill environment, as compared to free locomotion over solid and even ground, can reweight the sensory feedback due to many factors, such as the level of familiarity with the device, the dimensions of the belt or the stationarity of visual feedback (Van Hooren et al., 2019). The constraints imposed by the limited space and necessity of matching the belt’s speed (Dingwell and Cusumano, 2000), can act as external perturbations. However, our current results exclude that the CNS coped with those potential perturbations by widening the motor primitives.

Yet, when looking into the timing of main activation as described by the CoA, we found that motor primitives for treadmill locomotion were shifted earlier in time in both walking and running. This happened in one synergy out of four: the one for propulsion. The coordinated activity of foot plantarflexors characterize this synergy providing the main support and forward acceleration of the body mass (Arampatzis et al., 1999; Liu et al., 2008; Hamner and Delp, 2013; Santuz et al., 2018b; Bohm et al., 2019). It has been shown in humans that proprioceptive feedback from group II (muscle spindles) and/or group Ib (Golgi tendon organs) afferents is of paramount importance for the activation of plantarflexors (Dietz et al., 1994; Sinkjær et al., 2000). Additionally, mouse studies reported a crucial role of the proprioceptive feedback from plantarflexors in regulating the amplitude of muscle activity at different speeds (Mayer et al., 2018). We reinforced those observations showing that genetically modified mice lacking muscle spindles undergo a redistribution of the motor modules for propulsion when compared to wild type (Santuz et al., 2019). Moreover, we found that mutants could not manage to modulate the timing of motor primitives when external perturbations were added to locomotion (Santuz et al., 2019). Thus, the shifted CoA of the propulsion motor primitive might indicate that treadmill locomotion likely induced alterations in the proprioceptive sensory feedback from foot plantarflexors (i.e. PL, GM, GL, and SO). Additionally, we and others found similar shifts of the propulsion primitive’s CoA in both wild type mice (Santuz et al., 2019) and healthy humans (Maclellan et al., 2014; Santuz et al., 2018b) undergoing external perturbations, suggesting from yet another perspective that treadmills might perturb locomotion in ways that were never discussed before.

### Conclusions

In this study, we used a novel combination of machine learning and fractal analysis to understand those neuromotor control features of overground and treadmill locomotion that were not grasped by previous literature. Specifically, we found time-related alterations of motor primitives, the basic activation patterns common to functionally-related muscle groups. First, the primitives for the propulsion phase of both walking and running showed their main activation earlier in treadmill than in overground. This is similar to what previously reported for perturbed locomotion as compared to unperturbed. Second, motor primitives were on average more regular in treadmill than in overground locomotion. A data-driven suggestion that treadmills might constrain the muscle activation patterns for the control of human locomotion.

## Materials and Methods

This study was reviewed and approved by the Ethics Committees of the Humboldt-Universität zu Berlin. All the participants gave written informed consent for the experimental procedure, in accordance with the Declaration of Helsinki.

### Experimental protocol

For the experimental protocol we recruited 30 healthy and regularly active young adults (15 females, height 173 ± 10 cm, body mass 68 ± 12 kg, age 28 ± 5 years, means ± standard deviation). None of them was using orthotic insoles, had any history of neuromuscular or musculoskeletal impairments, or any head or spine injury at the time of the measurements or in the previous six months. All the volunteers completed a self-selected warm-up running on a treadmill, typically lasting 3 to 5 min (Santuz et al., 2016, 2018c). After being instructed about the protocol, they completed the measurements described in detail below.

The experimental protocol consisted of walking at 1.4 m/s and running at 2.8 m/s overground and on a single-belt treadmill (mercury, H-p-cosmos Sports & Medical GmbH, Nussdorf, Germany) equipped with a pressure plate recording the plantar pressure distribution at 120 Hz (FDM-THM-S, zebris Medical GmbH, Isny im Allgäu, Germany). The speeds were chosen since walking at 1.4 m/s and running at 2.8 m/s are close to the average comfortable locomotion speeds commonly reported in the scientific literature (Santuz et al., 2016, 2017a). Three pairs of photocells installed at 300 cm from each other were used to control the overground locomotion speed. After an accommodation period which usually involved 10 to 20 attempts to meet the requested speed, we recorded those trials that presented an error in matching the speed lower than ± 0.05 m/s in walking and ± 0.10 m/s in running.

### EMG recordings

The muscle activity of the following 13 ipsilateral (right side) muscles was recorded: *gluteus medius* (ME), *gluteus maximus* (MA), *tensor fasciæ latæ* (FL), *rectus femoris* (RF), *vastus medialis* (VM), *vastus lateralis* (VL), *semitendinosus* (ST), *biceps femoris* (long head, BF), *tibialis anterior* (TA), *peroneus longus* (PL), *gastrocnemius medialis* (GM), *gastrocnemius lateralis* (GL) and *soleus* (SO). The electrodes were positioned as previously reported (Santuz et al., 2018c, 2019). For the treadmill recordings, after the warm-up the participants were allowed to at least 60 s habituation (Santuz et al., 2018b). We recorded 10 overground and one treadmill trials (60 s) per locomotion type, per participant by means of a 16-channel wireless bipolar EMG system (Wave Plus wireless EMG with PicoEMG transmitters, Cometa srl, Bareggio, Italy) with an acquisition frequency of 2 kHz. For the recordings, we used foam-hydrogel electrodes with snap connector (H124SG, Medtronic plc, Dublin, Ireland). The overground trials were then concatenated (i.e. joined together) in a single one, so that for each participant four total trials were used for subsequent analysis: 1) concatenated overground walking; 2) concatenated overground running; 3) treadmill walking; 4) treadmill running. The first 30 gait cycles of each trial were considered for subsequent analysis (Santuz et al., 2018c). All the recordings can be downloaded from the supplementary data set, which is accessible at Zenodo (DOI: 10.5281/zenodo.3932767).

### Gait parameters

The gait cycle breakdown (foot touchdown and lift-off timing) was obtained by processing 3D acceleration data for the overground and plantar pressure distribution for the treadmill trials. For the segmentation of the overground attempts, we positioned a 3D accelerometer over the second-last pair of shoe eyelets, tightening the sensor using Velcro straps. We processed the obtained data using validated algorithms previously reported (Santuz et al., 2018b, 2020a). Treadmill recordings were segmented applying a previously published algorithm (Santuz et al., 2016) to the plantar pressure distribution data, recorded through the plate integrated in the treadmill. Other calculated gait spatiotemporal parameters were: stance and swing times, cadence (i.e. number of steps per minute), and the strike index, calculated as the distance from the heel to the center of pressure at impact normalized with respect to total foot length (Santuz et al., 2016). For stance, swing and cadence, we calculated the step-to-step percent coefficient of variation as the ratio between the standard deviation and the mean of each trial (Erra et al., 2019).

While for the treadmill trials the strike index was calculated by processing plantar pressure distribution data (Santuz et al., 2016), for the overground trials we made use of kinetics and kinematics data. In order to locate the center of pressure at touchdown, an infrared motion capture system (Vicon Nexus, version 1.7.1, Vicon Motion Systems, Oxford, UK) and a force plate (AMTI BP600, Advanced Mechanical Technology, Inc., Watertown, MA, USA) were used. Nine infrared cameras operating at 250 Hz recorded the position of two spherical reflective markers (ø 14 mm) placed on the heel and toe cap of the right shoe, approximately over the Achilles tendon insertion on the calcaneus and the first toe tip, respectively. The ground reaction forces were recorded at 1 kHz, and the center of pressure location during the stance phase was calculated using the recorded data. The participants were asked to walk or run on the straight pathway and were not told about the existence of the force plate. It was the operator’s task to check whether the plate was met by the right foot. If not, the trial was repeated. Strike index values range from 0 to 1, denoting the most posterior and the most anterior point of the shoe, respectively (Santuz et al., 2017b). Values from 0.000 to 0.333 are indication of a rearfoot strike pattern, while values from 0.334 to 1.000 represent a mid/forefoot strike pattern (Santuz et al., 2016).

### Muscle synergies extraction

Muscle synergies data were extracted from the recorded EMG activity through a custom script (R v3.6.3, R Core Team, 2020, R Foundation for Statistical Computing, Vienna, Austria) using the classical Gaussian NMF algorithm (Lee and Seung, 1999; Santuz et al., 2017a, 2018b, 2018c). The raw EMG signals were band-pass filtered within the acquisition device (cut-off frequencies 10 and 500 Hz). Then the signals were high-pass filtered, full-wave rectified and lastly low-pass filtered using a 4^th^ order IIR Butterworth zero-phase filter with cut-off frequencies 50 Hz (high-pass) and 20 Hz (low-pass for creating the linear envelope of the signal), as previously described (Santuz et al., 2018b).

After subtracting the minimum, the amplitude of the EMG recordings obtained from the single trials was normalized to the maximum activation recorded for every individual muscle. In other words, every EMG channel was normalized to its maximum for every trial (Santuz et al., 2018c, 2019, 2020a). Each gait cycle was then time-normalized to 200 points, assigning 100 points to the stance and 100 points to the swing phase (Santuz et al., 2017b, 2018b, 2018c, 2019, 2020a). The reason for this choice is twofold (Santuz et al., 2018c). First, dividing the gait cycle into two macro-phases helps the reader understanding the temporal contribution of the different synergies, diversifying between stance and swing. Second, normalizing the duration of stance and swing to the same number of points for all participants (and for all the recorded gait cycles of each participant) makes the interpretation of the results independent from the absolute duration of the gait events.

Synergies were then extracted through NMF as follows (Santuz et al., 2018b, 2018c). The 13 muscles listed above were considered for the analysis, (ME, MA, FL, RF, VM, VL, ST, BF, TA, PL, GM, GL and SO). The m = 13 time-dependent muscle activity vectors were grouped in a matrix V with dimensions m × n (m rows and n columns). The dimension n represented the number of normalized time points (i.e. 200*number of gait cycles). The matrix V was factorized using NMF so that V ≈ V_R_ = MP. The new matrix V_R_, reconstructed by multiplying the two matrices M and P, approximates the original matrix V. The motor modules (Gizzi et al., 2011; Santuz et al., 2017a) matrix M, with dimensions m × r, contained the time-invariant muscle weightings, which describe the relative contribution of muscles within a specific synergy (a weight was assigned to each muscle for every synergy). The motor primitives (Dominici et al., 2011; Santuz et al., 2017a) matrix P contained the time-dependent coefficients of the factorization with dimensions r × n, where the number of rows r represents the minimum number of synergies necessary to satisfactorily reconstruct the original set of signals V. M and P described the synergies necessary to accomplish the required task (i.e. walking or running, overground or on a treadmill). The update rules for M and P are presented in Equation (EQ1) and Equation (EQ2).

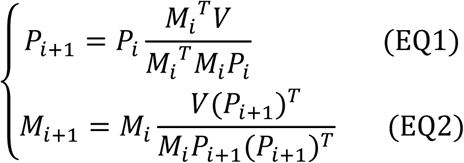

The quality of reconstruction was assessed by measuring the coefficient of determination R^2^ between the original and the reconstructed data (V and V_R_, respectively). The limit of convergence for each synergy was reached when a change in the calculated R^2^ was smaller than the 0.01% in the last 20 iterations (Santuz et al., 2017a) meaning that, with that amount of synergies, the signal could not be reconstructed any better. This operation was first completed by setting the number of synergies to one. Then, it was repeated by increasing the number of synergies each time, until a maximum of 10 synergies. The number 10 was chosen to be lower than the number of muscles, since extracting a number of synergies equal to the number of measured EMG activities would not reduce the dimensionality of the data. Specifically, 10 is the rounded 75% of 13, which was the number of considered muscles (Santuz et al., 2019). For each synergy, the factorization was repeated 10 times, each time creating new randomized initial matrices M and P, in order to avoid local minima (D’Avella and Bizzi, 2005). The solution with the highest R^2^ was then selected for each of the 10 synergies. To choose the minimum number of synergies required to represent the original signals, the curve of R^2^ values versus synergies was fitted using a simple linear regression model, using all 10 synergies. The mean squared error (Cheung et al., 2005) between the curve and the linear interpolation was then calculated. Afterwards, the first point in the R^2^-vs.-synergies curve was removed and the error between this new curve and its new linear interpolation was calculated. The operation was repeated until only two points were left on the curve or until the mean squared error fell below 10^−4^. This was done to search for the most linear part of the R^2^-versus-synergies curve, assuming that in this section the reconstruction quality could not increase considerably when adding more synergies to the model.

### Motor primitive geometrics and functional classification of synergies

We compared motor primitives by evaluating the one-dimensional statistical parametric mapping (SPM), center of activity (CoA), and the full width at half maximum (FWHM) (Cappellini et al., 2006, 2016; Pataky, 2010, 2012; Martino et al., 2014; Santuz et al., 2018b, 2019). The CoA was defined cycle-by-cycle as the angle of the vector (in polar coordinates) that points to the center of mass of that circular distribution (Cappellini et al., 2016), and then averaged. The polar direction represented the gait cycle’s phase, with angle 0 ≤ *θ*_*t*_ ≤ 2π. The following equations define the CoA:

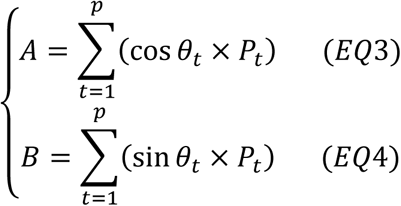

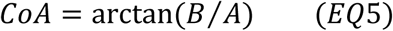

where *p* is the number of points of each gait cycle (*p* = 200). The FWHM was calculated cycle-by-cycle after subtracting the cycle’s minimum as the number of points exceeding each cycle’s half maximum, and then averaged (Martino et al., 2014). As a tool for visualizing differences in FWHM, we created heat maps (Figure 6) by counting cycle-by-cycle how many points of the relevant motor primitive exceeded half maximum and then averaging the obtained values over the 30 gait cycles per trial. The CoA and FWHM calculations and the subsequent SPM analysis were conducted only for the motor primitives relative to fundamental synergies. A fundamental synergy can be defined as an activation pattern whose motor primitive shows a single main peak of activation (Santuz et al., 2018b). When two or more fundamental synergies are blended into one (or when one synergy is split into one or more synergies), a combined synergy appears.

Combined synergies usually constitute, in our locomotion data, 10 to 30% of the total extracted synergies. While fundamental synergies can be compared given their similar function (i.e. motor primitives and motor modules are comparable since they serve a specific task within the gait cycle), combined synergies are often so different one from another that their classification is not possible. Due to the lack of consent in the literature on how to interpret them, we excluded the combined synergies from the FWHM analysis. The recognition of fundamental synergies was carried out by clustering similar motor primitives through NMF, using the same algorithm employed for synergy extraction with the maximum number of synergies set to the maximum factorization rank plus one. The obtained “principal shapes” for each of the four locomotion conditions were then compared to the motor primitives in order to cluster similar shapes. A primitive was considered similar to one of the principal shapes if the NMF weight was equal at least to the average of all weights. We then calculated the R^2^ of all the primitives that satisfied this condition, with the relevant principal shape. If the R^2^ was at least the 25% (or four times if the R^2^ was negative) of the average R^2^ obtained by comparing all the remaining primitives with their own principal shape, we confirmed the synergy as fundamental and classified it based on function. Primitives that were not clustered, were labeled as combined.

### Fractal analysis of motor primitives

To estimate the long-range dependence of motor primitives, we conducted a fractal analysis and calculated the Hurst exponent (H) following the rescaled range (R/S) approach (Hurst, 1951; Mandelbrot and Wallis, 1969). We proceeded as follows: 1) calculated the mean of the considered motor primitive of length n; 2) subtracted the mean to the original primitive; 3) calculated the cumulative sum of the obtained time series; 4) found the range R of this series (the range is the difference between the maximum and minimum values of a series); 5) calculated the standard deviation S; 6) computed the rescaled range R/S; 7) repeated the previous for N = n/2, n/4, n/8… and until a minimum of N=200, which is the normalized period of each motor primitive (Santuz and Akay, 2020); 7) calculated H as the slope of the log(N) vs. log(R/S) curve.

H can vary between 0 and 1. For 0.5 < H < 1, in the long term high values in the series will be probably followed by other high values (i.e. positive autocorrelation); in other words, the series is persistent or has long-term memory (Mandelbrot, 1983; Gneiting and Schlather, 2004; Tarnopolski, 2016). For 0 < H < 0.5, in the long term high values in the series will be probably followed by low values, with a frequent switch between high and low values (i.e. negative autocorrelation); in other words, the series is anti-persistent or has short-term memory (Mandelbrot, 1983; Gneiting and Schlather, 2004; Tarnopolski, 2016). A Hurst exponent of 0.5 indicates a completely random series without any persistence (Mandelbrot, 1983; Qian and Rasheed, 2004; Tarnopolski, 2016).

### Statistics

To investigate the effect of locomotion environment and type on the factorization rank, gait parameters, CoA, FWHM, and H of motor primitives and motor modules, we fitted the data using a generalized linear model with Gaussian error distribution. The homogeneity of variances was tested using the Levene’s test. If the residuals were normally distributed, we carried out a two-way repeated measures ANOVA with type II sum of squares for the dependent variables factorization rank, cadence, stance and swing time, CoA, FWHM, H, and muscle, the independent variables being the locomotion type (i.e. walking or running) and environment (i.e. overground or treadmill). If the normality assumptions on the residuals were not met, we used the function “raov”, a robust (rank-based) ANOVA from the R package Rfit (Kloke and McKean, 2012; McKean and Kloke, 2014). We then performed a least significant difference *post-hoc* analysis with false discovery rate adjustment of the *p*-values. Moreover, differences in motor primitives were tested using a two-way repeated measure ANOVA one-dimensional SPM (Pataky, 2010, 2012), with independent variables locomotion environment (i.e. overground or treadmill) and gait cycle (30 levels, each being one of the 30 recorded gait cycles per trial, per participant). To account for the bias related to the order of gait cycles, we performed the repeated measures ANOVA SPM 10000 times, randomizing at each repetition the gait cycle order for each participant. Results are reported as mean of the 10000 resamples.

All the significance levels were set to α = 0.05 and the statistical analyses were conducted using custom R v3.6.3 or Python (v3.8.2, Python Software Foundation, 2020, Wilmington, Delaware, United States) scripts. The spm1d (Pataky, 2012) open-source Python package v0.4.3 (spm1d.org) was used to generate F-values maps, F* limit and areas for the SPM analysis.

### Data availability

In the supplementary data set accessible at Zenodo (DOI: 10.5281/zenodo.3932767) we made available: a) the metadata with anonymized participant information; b) the raw EMG, already concatenated for the overground trials; c) the touchdown and lift-off timings of the recorded limb, d) the filtered and time-normalized EMG; e) the muscle synergies extracted via NMF; f) the code to process the data. In total, 120 trials from 30 participants are included in the supplementary data set.

The file “metadata.dat” is available in ASCII and RData format and contains:

- Code: the participant’s code
- Sex: the participant’s sex (M or F)
- Locomotion: the type of locomotion (W=walking, R=running)
- Environment: to distinguish between overground (O) and treadmill (T)
- Speed: the speed at which the recordings were conducted in [m/s] (1.4 m/s for walking, 2.8 m/s for running)
- Age: the participant’s age in years
- Height: the participant’s height in [cm]
- Mass: the participant’s body mass in [kg].

The files containing the gait cycle breakdown are available in RData format, in the file named “CYCLE_TIMES.RData”. The files are structured as data frames with 30 rows (one for each gait cycle) and two columns. The first column contains the touchdown incremental times in seconds. The second column contains the duration of each stance phase in seconds. Each trial is saved as an element of a single R list. Trials are named like “CYCLE_TIMES_P0020_TW_01,” where the characters “CYCLE_TIMES” indicate that the trial contains the gait cycle breakdown times, the characters “P0020” indicate the participant number (in this example the 20^th^), the characters “TW” indicate the locomotion type and environment (O=overground, T=treadmill, W=walking, R=running), and the number “01” indicate the trial number. Please note that the running overground trials of participants P0001, P0007, P0008 and P0009 only contain 21, 29, 29 and 26 cycles, respectively.

The files containing the raw, filtered, and the normalized EMG data are available in RData format, in the files named “RAW_EMG.RData” and “FILT_EMG.RData”. The raw EMG files are structured as data frames with 30000 rows (one for each recorded data point) and 14 columns. The first column contains the incremental time in seconds. The remaining 13 columns contain the raw EMG data, named with muscle abbreviations that follow those reported above. Each trial is saved as an element of a single R list. Trials are named like “RAW_EMG_P0003_OR_01”, where the characters “RAW_EMG” indicate that the trial contains raw emg data, the characters “P0003” indicate the participant number (in this example the 3^rd^), the characters “OR” indicate the locomotion type and environment (see above), and the numbers “01” indicate the trial number. The filtered and time-normalized emg data is named, following the same rules, like “FILT_EMG_P0003_OR_01”.

The muscle synergies extracted from the filtered and normalized EMG data are available in RData format, in the file named “SYNS.RData”. Each element of this R list represents one trial and contains the factorization rank (list element named “synsR2”), the motor modules (list element named “M”), the motor primitives (list element named “P”), the reconstructed EMG (list element named “Vr”), the number of iterations needed by the NMF algorithm to converge (list element named “iterations”), and the reconstruction quality measured as the coefficient of determination (list element named “R2”). The motor modules and motor primitives are presented as direct output of the factorization and not in any functional order. Motor modules are data frames with 13 rows (number of recorded muscles) and a number of columns equal to the number of synergies (which might differ from trial to trial). The rows, named with muscle abbreviations that follow those reported above, contain the time-independent coefficients (motor modules M), one for each synergy and for each muscle. Motor primitives are data frames with 6000 rows and a number of columns equal to the number of synergies (which might differ from trial to trial) plus one. The rows contain the time-dependent coefficients (motor primitives P), one column for each synergy plus the time points (columns are named e.g. “time, Syn1, Syn2, Syn3”, where “Syn” is the abbreviation for “synergy”). Each gait cycle contains 200 data points, 100 for the stance and 100 for the swing phase which, multiplied by the 30 recorded cycles, result in 6000 data points distributed in as many rows. This output is transposed as compared to the one discussed in the methods section to improve user readability. Trials are named like “SYNS_ P0012_OW_01”, where the characters “SYNS” indicate that the trial contains muscle synergy data, the characters “P0012” indicate the participant number (in this example the 12^th^), the characters “OW” indicate the locomotion type and environment (see above), and the numbers “01” indicate the trial number. Given the nature of the NMF algorithm for the extraction of muscle synergies, the supplementary data set might show non-significant differences as compared to the one used for obtaining the results of this paper.

All the code used for the pre-processing of EMG data and the extraction of muscle synergies is available in R format. Explanatory comments are profusely present throughout the script “muscle_synergies.R”.

## Acknowledgments

The authors are grateful to all the participants that showed great commitment and interest during the experiments and to Leon Brüll (Humboldt-Universiät zu Berlin) for the help with data organization.

## Author contributions

Conceptualization: E.P., A.A. and A.Sa.; Data curation: I.M., A.Se., V.M.M. and A.Sa.; Formal analysis: I.M. and A.Sa.; Investigation: I.M., A.Se., N.W., V.M.M., A.E. and A.Sa.; Methodology: A.A. and A.Sa.; Project administration: A.A. and A.Sa.; Resources: E.P., A.A. and A.Sa.; Software: I.M. and A.Sa.; Supervision: E.P., A.A. and A.Sa.; Validation: I.M., A.Se. and A.Sa.; Visualization: I.M. and A.Sa.; Writing – original draft: I.M., A.E., A.A. and A.Sa.; Writing – review & editing: I.M., A.Se., N.W., V.M.M., A.E., E.P., A.A. and A.Sa.

